# Two-point optical manipulation reveals mechanosensitive remodeling of cell-cell contacts in vivo

**DOI:** 10.1101/2022.07.08.499278

**Authors:** Kenji Nishizawa, Shao-Zhen Lin, Claire Chardès, Jean-François Rupprecht, Pierre-François Lenne

## Abstract

Biological tissues acquire reproducible shapes during development through dynamic cell behaviors. These events involve the remodeling of cell contacts driven by active cytoskeletal contractile forces. However how cell-cell contacts remodel remains poorly understood because of lack of tools to directly apply forces at cell-cell contacts to produce their remodeling. Here we develop a dual-optical trap manipulation method to impose different force patterns on cell-cell contacts in the early epithelium of the *Drosophila* embryo. Through different push and pull manipulations at the edges of junctions, the technique allows us to produce junction extension and junction shrinkage. We use these observations to constrain and specify vertex-based models of tissue mechanics, incorporating negative and positive mechanosensitive feedback depending on the type of remodeling. We show that Myosin-II activity responds to junction strain rate and facilitates full junction shrinkage. Altogether our work provides insight into how stress produces efficient deformation of cell-cell contacts in vivo and identifies unanticipated mechanosensitive features of their remodeling.

**Significance statement:** The highly organized tissues and organs that form our body emerge from internal dynamic activities at the cellular level. Among such activities, cell shape changes and cell rearrangement, cell extrusion and cell division sculpt epithelial tissues into elongated sheets, tubes and spherical cavities. Remodeling of cell-cell contacts, powered by actomyosin contractility, is key to all these transformations. Although much is known about the molecular machinery and biochemical signals that regulate remodeling of cell contacts, there is a lack of approaches to directly probe the mechanics of cell contacts and therefore assess their ability to resist or deform in response to mechanical loads. We developed an experimental technique to manipulate and exert contractile and extensile forces to cell-cell junctions. Our results lead to a specific physical model of junctional mechanics, with implications in the modeling of collective cell behavior in epithelial tissues.

## Introduction

How tissues and organs acquire their shape from internal cellular activities is a long-standing question that has fascinated several generations of scientists. Over the past decades, fluorescence imaging has revealed the complex choreography of cells during tissue morphogenesis (1, 2). By integrating imaging with genetics and biochemical manipulations, several studies have identified the molecular players shaping tissues and how their activities are regulated (3–6). Cell shape changes and cell rearrangements are among the most prominent cellular behaviors of tissue morphogenesis, enabling epithelial tissues to flow, elongate, fold or invaginate (7). Such behaviors involve constant gain and loss of cell-cell contacts, through remodeling of cell-cell junctions, adherens junctions in particular (8).

Recent attempts to understand the mechanics of cell junctions have highlighted how the interplay between actomyosin contractility and adhesion affects the length of junctions (9–11) but also their dissipative mechanical nature and their mechanosensitive response (12–15). However, it remains to be understood how the distribution of forces at the cell and tissue scales produce their remodeling.

The theoretical framework of tissue mechanics, notably the numerical schemes called vertex models (16, 17) are designed to bridge the mechanics of cell junctions to tissue organization. Vertex models have been applied to different morphogenetic movements such as epithelial folding (18) and extension (19), formation of tissue boundaries (20), of the optical cup (21) or of the regular patterns in the retina (22). However, they rely on assumptions of junction mechanics that have been poorly assessed, limiting their ability to faithfully predict collective cell behaviors.

Here we implement a method based on optical manipulation that allows us to directly remodel cell junctions and thus discriminate between different models of epithelial mechanics. The optical method is derived from our previous work where we showed that optical tweezers can directly trap individual junctions and locally deform them (23). Extending the method to deflect two junctions, we apply different patterns of forces at the edges of junctions to produce different modes of remodeling, including extension and shrinkage (partial or total). We used these measurements to challenge recently proposed models of junctional remodeling (14, 24). Going back and forth between experiments and simulations, we specify the mechanosensitive response of the junctions in vivo.

## Results

### Junction extension

To probe the mechanics of remodeling of cell junctions, we developed an experimental approach that uses two optical traps (Fig. 1A and 1B). The experimental setup combines a home-built optical tweezers system with a spinning disk microscope for high-resolution imaging (Fig. 1A). The two traps are generated and controlled by fast galvanometric mirrors that time-share an infrared laser beam between two positions. We have shown previously that a single optical trap can directly tweeze and deflect individual cell junctions, enabling the measurement of junction tension and junction rheology in the early epithelium of the *Drosophila* embryo (12, 23). However, single junction manipulation does not result in significant contact remodeling. We reasoned that by tweezing and manipulating two junctions concomitantly, we could deform their adjacent common junction (Fig. 1B, black junction in the sketch). We first applied this strategy in a diagonal pull configuration, where the two traps are moved away from each other in an antiparallel direction by 1.5 μm, then maintained at fixed positions (Fig. 1B and Movie 1). This manipulation results in the extension of the common adjacent junction, called hereafter the middle junction (Fig. 1C, length change of the black junction sketched in Fig. 1B). The middle junction extends gradually and reaches a stable length within 60 s (ΔI_60s_, Fig. 1C), which corresponds to a time slightly larger than the timescale of mechanical dissipation that we measured previously in this early epithelium (50 s see (12)). The diagonal pull with two traps results in asymmetric length changes of the adjacent junctions (Fig. 1D). The manipulated junctions (Fig. 1D, red) are first deflected but do not change their length over the long term. In contrast, the adjacent non-manipulated junctions decrease their length (Fig. 1D, blue). We checked that the observed length changes were not a consequence of changes in z-axis of the cells due to trapping forces (Supplementary Fig. S1).

**Figure 1.**
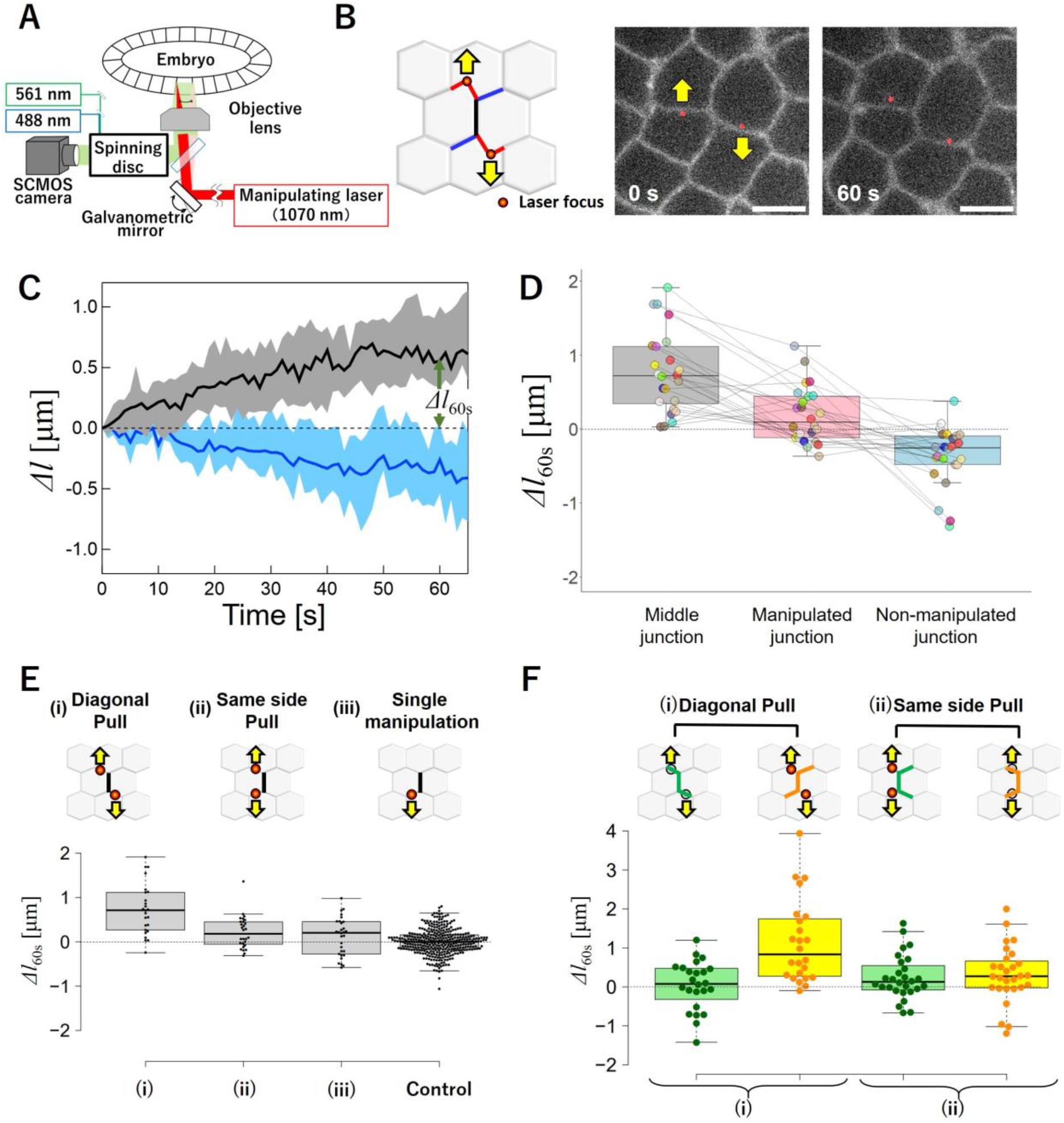
Remodeling cell contacts by two-point optical manipulation. (A) Schematic of the setup. The early epithelium of the *Drosophila* embryo is imaged using a spinning disc confocal system while cell-cell junctions are remodeled by a dual optical trap produced by an infrared laser. (B) Schematic (left) and images (right) of *diagonal pull* manipulation: the two traps (red dots) are positioned at two junctions (red, *manipulated junctions*) sharing a common neighboring junction (black, *middle junction*). The traps are moved away from each other, in antiparallel direction and maintained at 1 μm distance from their initial position (yellow arrows), causing junction remodeling. The cell junctions are labelled by E-cad::GFP. Scale bar: 5 μm. (C) Length changes of the middle (black) and adjacent non-manipulated junctions (blue). Median (solid line) and the 25^th^-75^th^ percentile range are shown. ΔI_60s_ denotes the length change 60 s after the onset of manipulation (N=25). (D)Length change at 60 s for the different junctions. (E) Comparison of length changes at 6 0s of the middle junction for different manipulation geometries: (i) diagonal pull (N=25), (ii) same side pull (N=28), (iii) single junction pull (N=30). Control corresponds to length changes at 60 s of junctions in the absence of manipulation. (F) Comparison of cumulative length changes showing compensatory mechanism between extension of the middle and contraction of the adjacent non-manipulated junctions, f*or diagonal pull*.

To test if other patterns of forces can remodel cell junctions, we then manipulated two junctions belonging to the same cell and sharing an adjacent junction (configuration called *same side pull*, Fig. 1E (ii)). In contrast to the *diagonal pull* (Fig. 1E (i)), the *same side pull* manipulation did not produce any significant length changes of the middle junction (Fig. 1E (ii)), nor that of the manipulated or adjacent non-manipulated junctions (Fig. 1F (ii) to be compared to diagonal pull (i)). Note that single junction manipulation is also inefficient in changing the length of adjacent junctions (Fig. 1E(iii)). The diagonal pull elicits an asymmetric sliding mechanism (Fig. 1F (i) green versus orange and Fig. 1D): the middle junction extends at the expense of adjacent non-manipulated junctions while the manipulated junctions maintain their length.

To understand the mechanical origin of this response, we adapted a mechanical model of tissues that could be experimentally falsifiable. In vertex models, the geometry of epithelial cells is represented by edges (junctions) connecting vertices (representing tricellular contacts) at a set of location ***r**_i_*. The motion of the vertices is then determined by the balance between friction forces (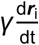, with *γ* a friction coefficient) and the mechanical work exerted by pressure and tension forces, described through a function *U* that includes an elastic control of the cell area (modeling the difference between cell pressure) and a set of junction tension (17):

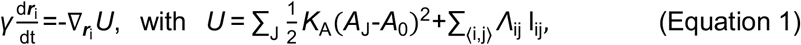

where *A*_J_ denotes the cell area of the cell J and *A*_0_ its target area; *K*_A_ the area elastic modulus; *Λ*_ij_ the junction tension and l_ij_ the length along the junction between the vertices i and j.

To model the deformations induced by the optical traps, we add a linear trapping force 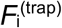 on the deflected vertices

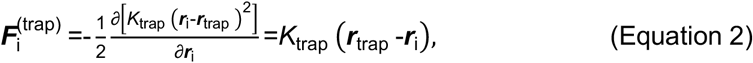

where *K*_trap_ is the trap stiffness and ***r***_tap_ and ***r***_i_ denote the trap and junction positions, respectively. For several manipulations, ***r***_i_ does not correspond to the location of a particular tri-cellular junction. We therefore considered a modified vertex model where we generated a set of two-way vertices, initiated at the middle of all junctions; defining such two-way vertices allows us to mimic the effect of the trapping forces, which, when active, induce junction bending (Fig. 2A and Movie 2).

**Figure 2.**
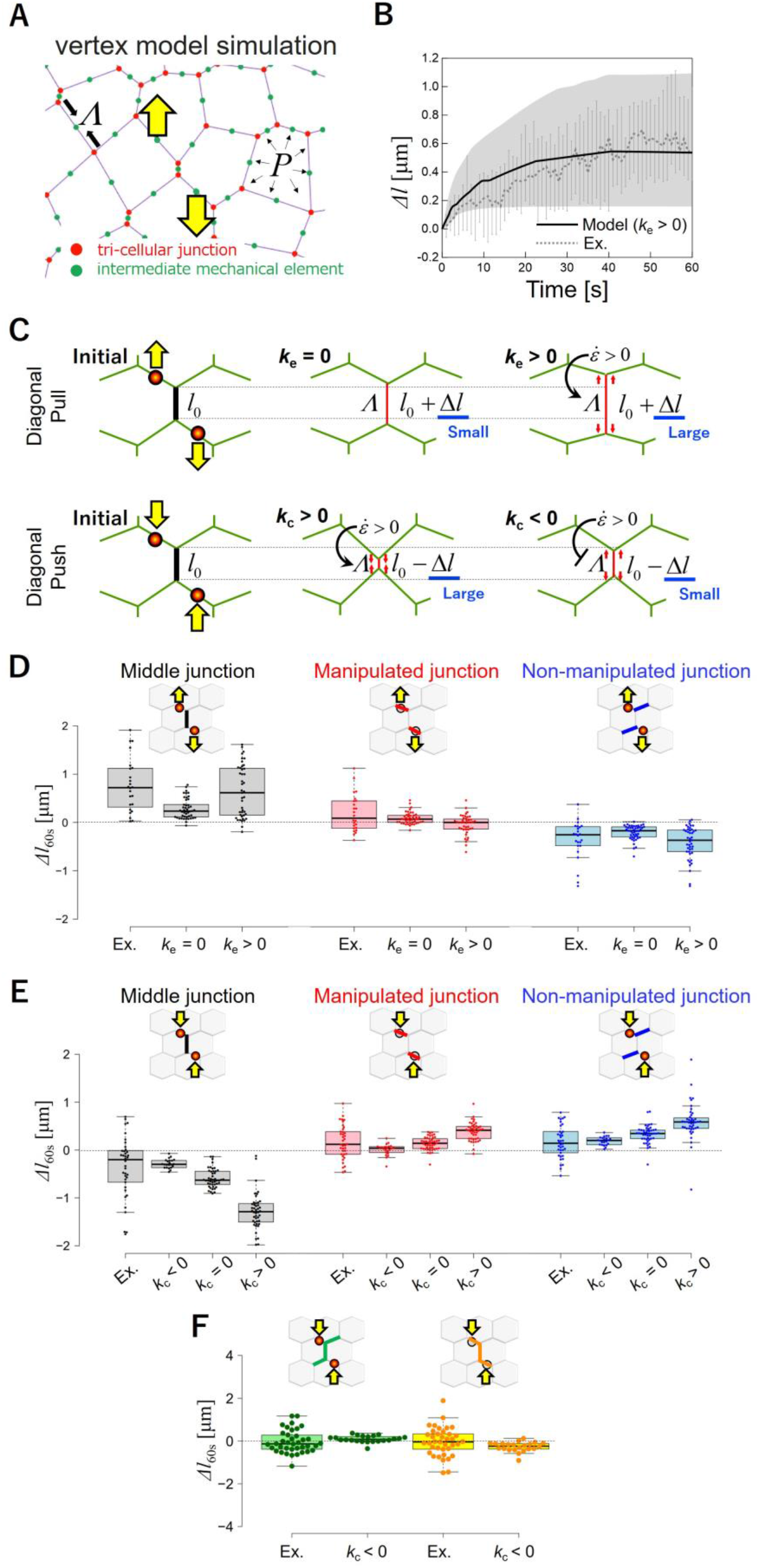
Vertex model simulations including dynamic feedback predict junction extension and contraction. (A) A vertex-based network with tricellular junctions (red dots) and intermediated mechanical elements (green dots), on which trapping forces are applied (yellow arrows). The model considers tensions *Λ*_ij_ along junctions and cell pressures within each cells *P* = *K*_A_(*A*-*A*_0_), with *A* (resp. *A*_0_) the cell current (resp. target) area (B) Model simulated (solid curve) and experimentally observed (broken curve) length changes of the middle junction (see Methods for details). (C) Predicted length changes for *diagonal pull* and *diagonal push* manipulation, in the presence or absence of dynamic remodeling. In absence of dynamic remodeling *k*_e_ *k*_e_ =0), extensile strain does not produce any change of junction tension Λ (middle top panel). Extensile strain 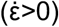 caused by *diagonal pull* elicits a negative feedback (*k*_e_>0) on tension, which favors larger extension (top right). Contractile strain 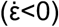 caused by *diagonal push* can elicit a positive feedback (*k*_c_>0, middle bottom) or a positive feedback (*k*_c_<0, right bottom) on tension, which favors large or small contraction, respectively. (D) Comparison of model simulated and experimentally observed length changes at 60s post-manipulation for middle, manipulated and non-manipulated adjacent junctions, for *diagonal pull* (N=25). (E) Comparison of model simulated and experimentally observed length changes at 60s post-manipulation for middle, manipulated and non-manipulated adjacent junctions, for *diagonal push* (N=38). (F) Comparison of simulated and experimentally observed cumulative length changes at 60s post-manipulation showing compensatory mechanism between contraction of the middle and extension of adjacent junctions, for *diagonal push*.

Combining such trap model with the energy Eq. (1) defines what we call our model A. We ran simulations on networks of cells whose statistical properties matched those observed in experiments in terms of area, perimeter, length of cell-cell edges, tension and pressure statistics (see Supplementary Materials). We find that such model A failed to reproduce the extension observed in *diagonal pull* manipulation (Fig. 2A). This indicates that a mechanism that favors large strains in response to extensile forces is missing in the model A.

We then adapted a vertex model adaptation in which junctions are dynamically remodeled upon large strain; following (14). We consider that the junction tension *Λ*_ij_ along the junction between the vertices i and j (Fig. 2C) dynamically adapts to changes in junction length according to the rule

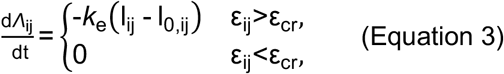

where *k*_e_ denotes the rate of tension remodeling under junction extension; ε_ij_= (I_ij_-I_0,ij_)/I_0,ij_) along the strain along the junction between the vertices i and j and ε_cr_ represents a critical strain beyond which such remodeling occurs. Following (14), the rest length I_0,ij_ of the cell-cell junction evolves according to

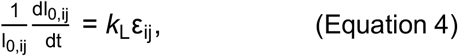

where *k*_L_ > 0 is a viscous relaxation rate (note that at steady state, I_ij_=I_0,ij_).

Adding this mechanosensitive response to the model A defines what we call the model B (model A and B are identical when *k*_e_ = 0). For a positive value *k*_e_ = 0.005 (= 0.18 pN·μm^−1^·s^−1^) together with *k*_L_ = 0.04 (=0.04 s^−1^), we accurately predict the experimental observations (Fig. 2B). Such behavior reveals that an extensional strain reduces tension (Fig. 2C). The model B also accurately predicted the reduction in length of the adjacent non-manipulated junctions (Fig. 2D). The model B further predicted the fact that, in the *same side pull* manipulation (Table 1), the junction length changes would be limited.

**Table 1.**
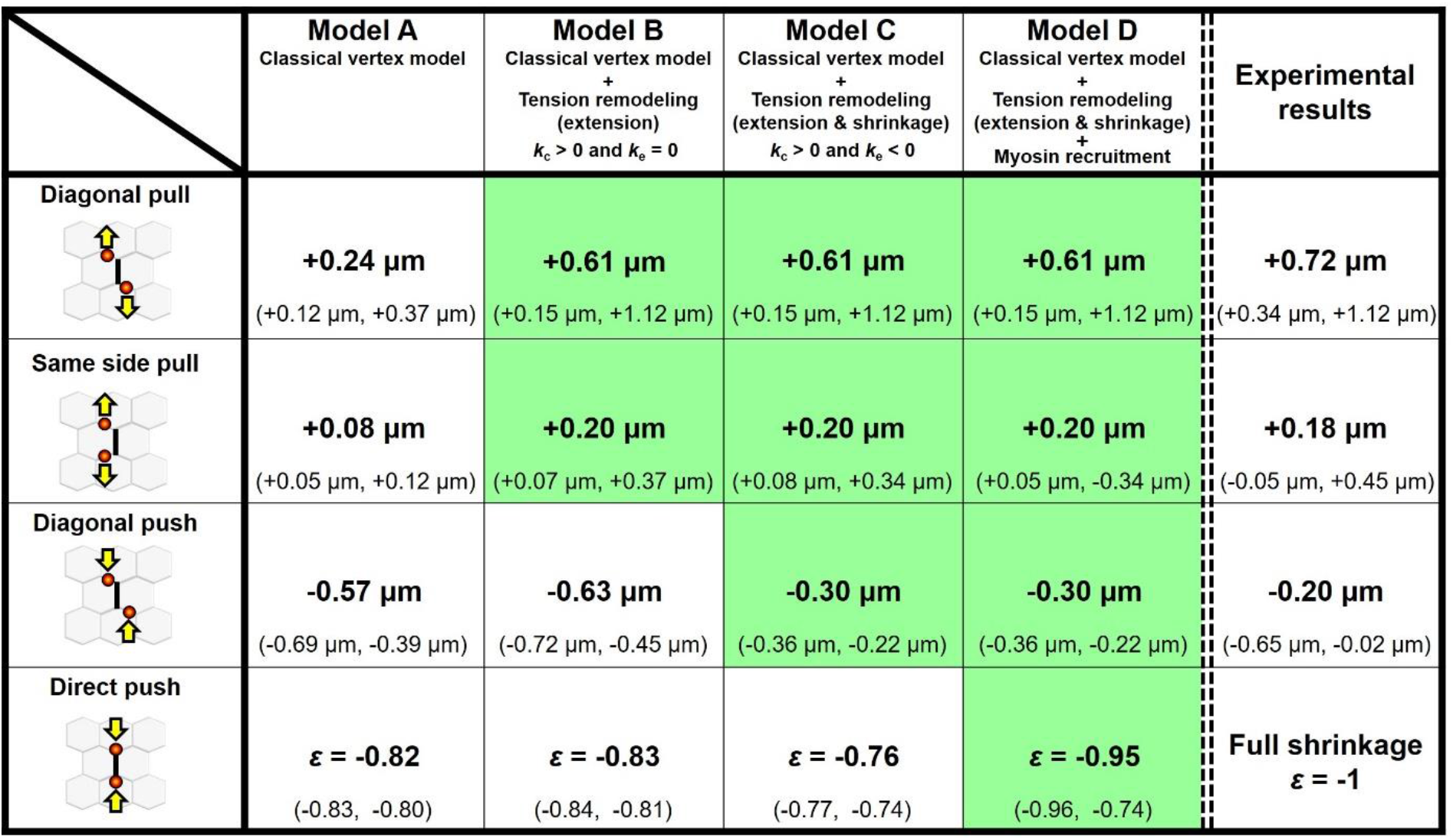
Comparison between experimental observations and models. Comparison of the extension of middle junction (diagonal pull, same side pull, and diagonal push) or the strain of middle junction (direct push) between experimental observations and models for the different types of manipulation. Medians and 25th-75th percentiles. Model predictions that are closest to the experimental data are highlighted in green.

| | **Model A**<br>Classical vertex model | **Model B**<br>Classical vertex model<br>+<br>Tension remodeling<br>(extension)<br>$k_c > 0$ and $k_e = 0$ | **Model C**<br>Classical vertex model<br>+<br>Tension remodeling<br>(extension & shrinkage)<br>$k_c > 0$ and $k_e < 0$ | **Model D**<br>Classical vertex model<br>+<br>Tension remodeling<br>(extension & shrinkage)<br>+<br>Myosin recruitment | **Experimental**<br>**results** |
| --- | --- | --- | --- | --- | --- |
| **Diagonal pull** | **+0.24 $\mu\text{m}$**<br>(+0.12 $\mu\text{m}$, +0.37 $\mu\text{m}$) | **+0.61 $\mu\text{m}$**<br>(+0.15 $\mu\text{m}$, +1.12 $\mu\text{m}$) | **+0.61 $\mu\text{m}$**<br>(+0.15 $\mu\text{m}$, +1.12 $\mu\text{m}$) | **+0.61 $\mu\text{m}$**<br>(+0.15 $\mu\text{m}$, +1.12 $\mu\text{m}$) | **+0.72 $\mu\text{m}$**<br>(+0.34 $\mu\text{m}$, +1.12 $\mu\text{m}$) |
| **Same side pull** | **+0.08 $\mu\text{m}$**<br>(+0.05 $\mu\text{m}$, +0.12 $\mu\text{m}$) | **+0.20 $\mu\text{m}$**<br>(+0.07 $\mu\text{m}$, +0.37 $\mu\text{m}$) | **+0.20 $\mu\text{m}$**<br>(+0.08 $\mu\text{m}$, +0.34 $\mu\text{m}$) | **+0.20 $\mu\text{m}$**<br>(+0.05 $\mu\text{m}$, -0.34 $\mu\text{m}$) | **+0.18 $\mu\text{m}$**<br>(-0.05 $\mu\text{m}$, +0.45 $\mu\text{m}$) |
| **Diagonal push** | **-0.57 $\mu\text{m}$**<br>(-0.69 $\mu\text{m}$, -0.39 $\mu\text{m}$) | **-0.63 $\mu\text{m}$**<br>(-0.72 $\mu\text{m}$, -0.45 $\mu\text{m}$) | **-0.30 $\mu\text{m}$**<br>(-0.36 $\mu\text{m}$, -0.22 $\mu\text{m}$) | **-0.30 $\mu\text{m}$**<br>(-0.36 $\mu\text{m}$, -0.22 $\mu\text{m}$) | **-0.20 $\mu\text{m}$**<br>(-0.65 $\mu\text{m}$, -0.02 $\mu\text{m}$) |
| **Direct push** | **$\varepsilon = -0.82$**<br>(-0.83, -0.80) | **$\varepsilon = -0.83$**<br>(-0.84, -0.81) | **$\varepsilon = -0.76$**<br>(-0.77, -0.74) | **$\varepsilon = -0.95$**<br>(-0.96, -0.74) | **Full shrinkage**<br>**$\varepsilon = -1$** |

### Junction shrinkage

As the *diagonal pull* manipulation produces significant extension of the middle junction, we wondered if the reverted manipulation (which we call *diagonal push*) could produce the reverse length change (Fig. 2C bottom and 2E). We found that such push manipulation leads to a significant shrinkage of the middle junction, and to an extension of the manipulated and non-manipulated adjacent junctions. In contrast to the *diagonal pull* manipulation, the *diagonal push* produces a symmetrical deformation (Fig. 2F): the shrinkage of the middle junction is similarly compensated by the extension of the manipulated and non-manipulated adjacent junctions. As for junction extension, we used our observations to constrain the physical model. We noticed that the extent of junction shrinkage in *diagonal push* was smaller than that for junction extension in *diagonal pull*. We thus tested if dynamical remodeling was also present or not for junction shrinkage by introducing an asymmetric junctional remodeling in the form

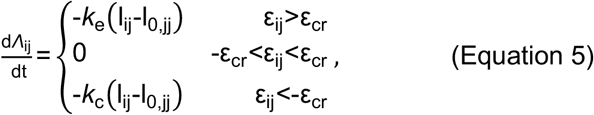

where we introduce the parameter *k*_c_ for the rate of tension remodeling under junction contraction. Exploring different values of *k*_c_ (Fig. 2C bottom, *k*_c_ = 0, *k*_c_ < 0, *k*_c_ > 0), we found that a negative value *k*_c_ =−0.003 (=−0.10 pN·μm^−1^·s^−1^, see Methods) produced the best fit to our experimental observations, not only for the shrinkage of the middle junction but also for manipulated and non-manipulated adjacent junctions (Fig. 2E). The fact that the best fit value of *k*_c_ is negative indicates that under this manipulation junction shrinkage reduces junction tension (Fig. 2C bottom). A prediction of this mechanosensitive response is that after trap release, the middle junction should relax back to a configuration associated with a lower tension. Consistent with our model prediction, we found that after trap release, the middle junction returns to a longer length (see Supplementary Fig. S2).

In 5 instances out of 38, the *diagonal push* manipulation led to a full shrinkage of the middle junction, i.e. with the two three-way vertices on each side of the junction joining into a single four-way vertex.

The formation of four-way vertices is a step in the cell-cell intercalation process observed in vivo (25). To gain understanding in the mechanics of such full junction remodeling, we designed the *direct push* manipulation, by positioning the traps on the two vertices of a junction and moving them inwards to the middle of the junction (Fig. 3A and 3B, and Movies 3 and 4). Such *direct push* manipulation led to full-junction shrinkage on a time scale that varied between a few seconds to more than 100 s (Fig. 3B, 3C and 3D). We compared these striking observations with the predictions of the model C elaborated so far. While such model C qualitatively explains the junction shrinkage of the targeted junction and the extension of its neighboring junctions, the model C predicts a partial shrinkage (at 80% of the initial length) and fails to explain full (100%) shrinkage of the middle junction (Fig. 3E and Table 1).

**Figure 3.**
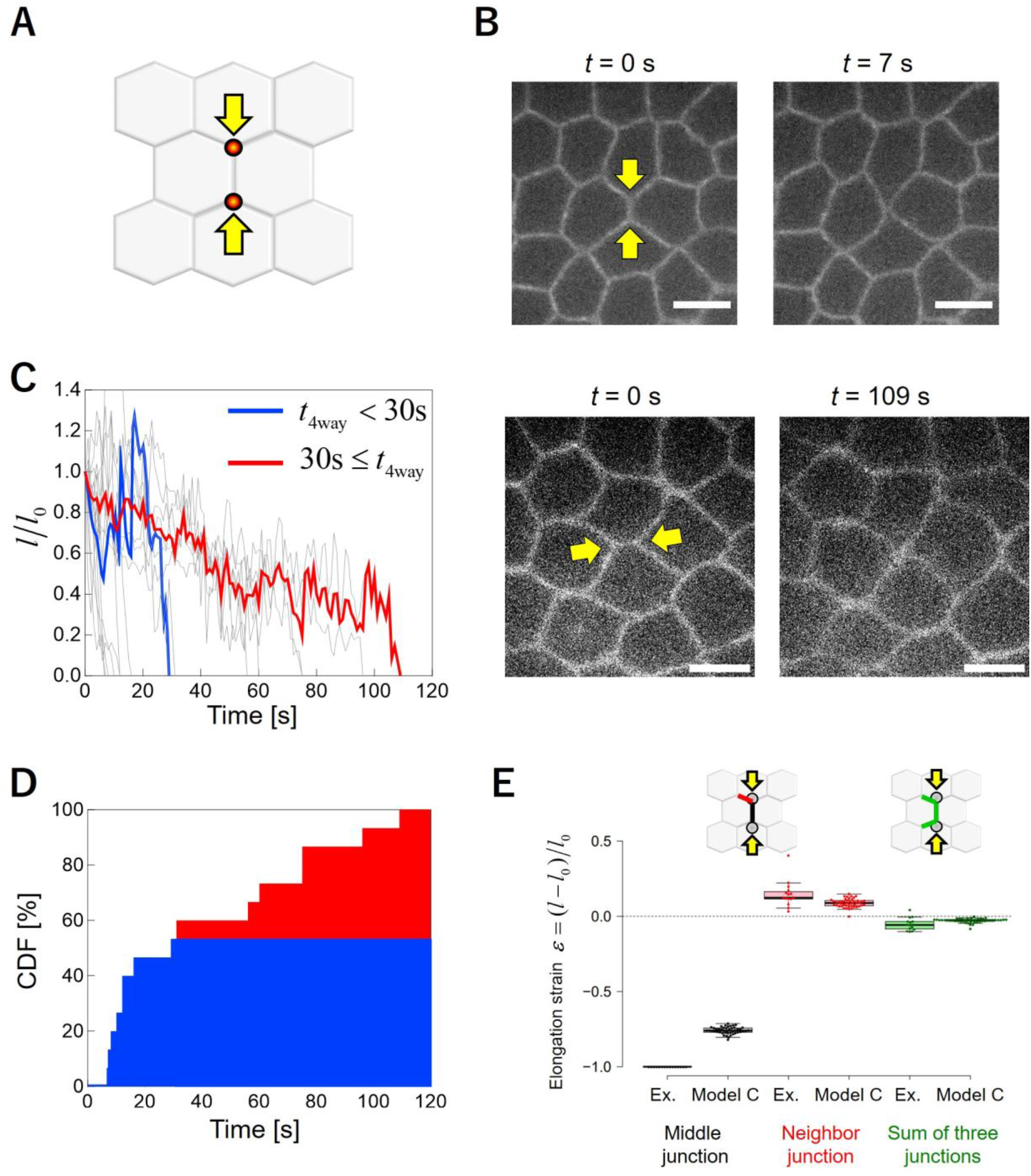
Direct push on vertices produces full junction remodeling. (A) Schematic of *direct push* manipulation. Traps are positioned on vertices and move towards the center of the middle of the junction (B) Two examples of full-junction shrinkage with distinct kinetics: fast (top image, 7 s) and slow (109 s) shrinkage. Scale bar: 5 μm. (C) Length changes of shrinking junctions over time (N=8). Full shrinkage times are broadly distributed. Blue and red curves show mean length changes for curves exhibiting short full shrinkage times (t_4way_<30 s) and long full shrinkage times (t_4way_≥30 s). Scale bar: 5 μm (D) Cumulative density function of full-junction shrinkage time (N=7). (E) Comparison of model simulated and experimentally observed strains. The model C does not predict full junction shrinkage.

### Myosin-II feedbacks on junction shrinkage

Such discrepancy in the *direct push* prediction points to a missing mechanism in our model that would positively feedback on the optically forced shrinkage. Here we find that Myosin-II (Myo-II) recruitment can mediate such mechanism. Myo-II is known to be required for cell intercalation and in particular for junction shrinkage during germband elongation in the early *Drosophila* epithelium (25). During this process Myo-II accumulates along shrinking junctions in a pulsatile fashion (4). As suggested in several works (26–30), we hypothesized that Myo-II could react to deformation. To test this hypothesis, we performed experiments in Rock-inhibited embryos (using the Rho-kinase inhibitor H-1152) and compared the deformations produced in this condition with untreated embryos, both for *direct push* and *diagonal pull* (Fig. 4A and 4B). We did not manage to shrink junctions by *direct push* of vertices in Rock-inhibited embryos (Fig. 4A, bottom images and Movie 5). Instead, the manipulation resulted in the junctions being bent (Movie 5). This shows that Myo-II activity is required for full-junction shrinkage process. In contrast, for diagonal pull, we did not observe any difference in junction deformation between the Rock-inhibited and untreated embryos (Fig. 4B, bottom plots). To further test the requirement of Myo-II in junction shrinkage in *push* experiments, we imaged Myo-II during optical manipulation (Fig. 4C and 4D). Analysis of Myo-II intensity along the junction shows the temporal accumulation of Myo-II in the targeted junction (kymograph Fig. 4D). This observation supports a Myo-II dependent feedback mechanism that would amplify the reduction in junction length by optical manipulation and contribute to full-junction shrinkage (Fig. 4E). To further assess this point, we looked for correlations between Myo-II intensity and the strain rate in *push* experiments (Fig. 4F and 4G). Optical manipulation allows us to explore a regime of high (negative) strain rates (circles and diamonds) compared to those observed in absence of manipulation (triangles), and reveals the mechanosensitive response of Myo-II to strain rate (Fig. 4G).

**Figure 4.**
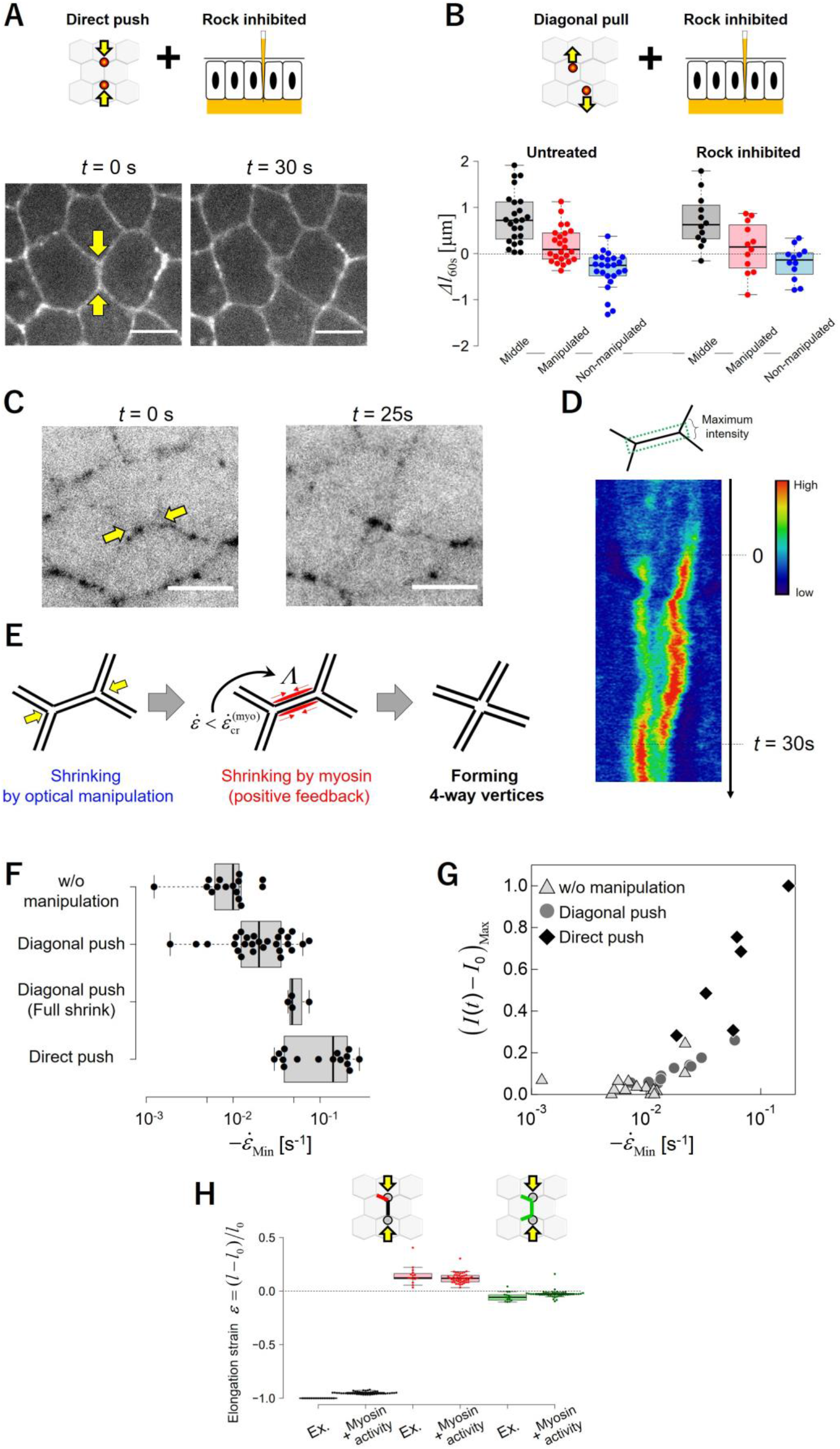
Myosin-II activity depends on contractile strain rate to produce full shrinkage. (A) *Direct push* manipulation in Rock-inhibited embryos (via injection). At 30 s post-manipulation, the middle junction appears bent but not shrunk. Scale bar: 5 μm. (B) *Diagonal pull* manipulation in Rock-inhibited (N=11) and comparison to untreated embryos (N=25). Length changes in the two conditions for middle (black), manipulated (red) and non-manipulated adjacent (blue) junctions. (C) Images of Myo-II (Sqh::GFP, inverted contrast) at the onset of the direct push and during shrinkage (25 s). Scale bar: 5 μm. (D) Kymograph of Myo-II (Sqh::GFP) intensity along a shrinking junction under *direct push*, showing accumulation of Myo-II over time. (E) Schematic of the mechanical model showing the feedback mechanism by which Myo-II strain rate dependent feedback on tension promotes full junction shrinkage. (F) The minimal strain rate (i.e. maximum absolute contraction rate), for *direct push* manipulation, *diagonal push* manipulation (making four-way vertices), *diagonal push* manipulation and in absence of manipulation. (G) Myo-II maximum intensity along junctions as a function of the minimal strain rate, for *direct push* manipulation (diamonds), diagonal push manipulation (circles), and in absence of manipulation (triangles). (H) Comparison of model D simulated and experimentally observed strains. The model D includes Myo-II positive feedback on tension depicted in (E).

This demonstrates the existence of a positive feedback mechanism that we introduced into our physical model. We assumed that below a critical strain rate 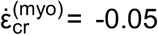 (−0.05 s^−1^, see Methods). Myo-II contractility would produce an active tension that writes

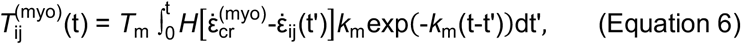

where *H*(*x*) is the Heaviside step function (equal to 1 if *x* > 0, 0 otherwise); 1/*k*_m_ is a time scale for myosin recruitment and *T*_m_ a strength of the myosin induced tension modulation. The addition of such relation to our previous model C defines model D. Using such model D, with *k*_m_ = 0.10 (= 0.10 s^−1^) and *T*_m_ = 0.25 (= 54 pN), we can recapitulate the full junction shrinkage observed in the *direct push* manipulation (Fig. 4H), while accounting for all other experimental observations (Table 1).

## Discussion

Here, we have introduced a method to efficiently remodel cell junctions. While single trap manipulation provided means to measure local junction tension and junction rheology (23, 31), we showed that dual trap manipulation can exert forces in several configurations to extend or shrink junctions. This allows not only to directly mimic events that occur in vivo, but also to assess, without proxy, the mechanics of junction remodeling.

By applying different patterns of forces, we identified different modes of deformation. These modes involve length compensatory mechanisms between adjacent junctions. Diagonal pull elicits junction extension by asymmetric sliding, as evidenced by the asymmetry of length changes of the non-manipulated and adjacent manipulated junctions (Fig. 1D and 1F). Such asymmetry suggests that the two apposed membranes forming the extending junction slide along each other. Whether the resulting shear could lead to the rupture of adhesion bonds remains to be understood (32). In contrast, shrinkage by *diagonal push* or *direct push* results in a symmetric sliding with equal compensation of length from the non-manipulated and adjacent manipulated junctions. This suggests an unzipping mechanism, in agreement with recent observations of vertex sliding mediated by Myo-II contractility (33).

In recent years, several techniques to measure tissue mechanics in situ have burgeoned (34–36), but with a few exceptions in vivo (12) and in vitro (14, 37), these are seldom used to falsify theoretical models. Here we have used dual-trap manipulation to discriminate among several mechanical models. We found that junction remodeling cannot be explained by the usual standard classical vertex models with constant tensions. We find that a set of simple mechanosensitive rules, implying positive and feedback loops on the junctional tension, allows us to describe our complete set of experimental manipulations. Our direct experimental approach complements image-based force inference methods, that have been recently implemented to estimate the mechanical parameters of remodeling epithelial tissues (38).

Our findings extend recent measurements of the viscoelastic and mechanosensitive properties of junctions. Single trap optical manipulation has revealed the characteristic time of energy dissipation at junctions, which is dependent on actin turnover (12). Optogenetic control of Myo-II activity has shown that the viscoelastic response of junctions is active: upon strain, junctions adapt their contractile tension. Here we specify the positive and negative feedbacks that regulate junction dynamics. Consistent with in vitro results (14), we found that junctional tension adapts in vivo to the applied strain. However, we found that contraction and extension lead to distinct adaptation. Furthermore, our approach disentangles the Myo-II dependent and Myo-II independent contributions to strains. On the one hand, we identified two Myo-II-independent mechanical feedbacks: one positive feedback to junction extension (as encapsulated in the sign of the parameter *k*_e_) and one negative feedback to junction contraction (*k*_c_ < 0). On the other hand, we uncovered a Myo-II dependent feedback to junction contraction (rate *k*_m_), that is controlled by the junctional strain-rate, complementing previous reports showing that Myo-II accumulates upon tissue deformation (26–30). We envisage that such set of negative/positive couplings between forces and deformation could represent an efficient mechanism to buffer weak contraction events, while ensuring full contraction at large loads. As such, we expect our findings to have broad implications for the understanding of tissue morphogenesis.

## Acknowledgments

The work is supported by the Agence Nationale de la Recherche (“Investissements d’Avenir”, ANR-16-CONV-0001 from Excellence Initiative of Aix-Marseille University -A*MIDEX, and generic grants, ANR-17-CE13-0032 and ANR-20-CE30-0023). We also acknowledge the France-Bioimaging Infrastructure (ANR-10-INBS-04). We thank the Lenne, Rupprecht and Lecuit groups for discussions.

## Methods

### Sample preparation

To image the adherens junctions and Myo-II in *Drosophila* embryos, we used E-cadherin::GFP flies (endogenous promoter) and sqsh::GFP flies, respectively. Once harvested, the embryos were washed with 100% bleach for 50 seconds to remove the chorion. Embryos at the end of cellularization (stage 5 end) were then selected under a dissection microscope and aligned on the edge of the coverslip. Alignment was done with the germband visible in the imaging plane. For Myo-II activity inhibition, embryos were placed in halocarbon oil and injected using a microinjection setup with ROCK inhibitor (H-1152, 40 mM, Invitrogen), Embryos were immersed in halocarbon oil for spinning disk imaging.

### Two-point optical manipulation and imaging

Optical manipulation of the cell junctions in individual embryos was done using a spinning-disk microscope (Perkin-Elmer), coupled with a home-built dual trap laser system. A 100x water immersion lens (Nikon) was used for imaging and optical manipulation in the imaging plane. To achieve two-point manipulation, we split an IR laser beam (1070 nm wavelength) into two beams by fast commutation (every 4 ms) of two galvanometric mirrors. The relationship between the (x,y) positions of the two resulting traps and the galvanometric command voltages is calibrated by trapping colloidal beads in water prior manipulation in the embryo (23).

For junction manipulation, laser traps are first stably positioned 5 s on the junctions, then moved to a defined distance and maintained (Fig. 1A). For most manipulation the trap displacements were 1 μm within 3 s and laser power per trap fixed at 200 mW.

### Image and data analysis

To analyze the length of junctions, we performed image segmentation with Tissue analyzer (39). The length changes presented in the figures are averaged over 5s. To measure the strain rate, the curves are first smoothed using a smoothing spline with a smoothing factor of ~5 (applied to the data acquired with 1 second frame rate).

smoothing spline). To analyze the intensity of Myo-II, we corrected for photobleaching (Fiji) and then measured the signal intensity at the junctions from the segmented image. Myo-II intensity is defined by the ratio of the integrated signal at the junction to the junction length. In Fig. 4, the increase in Myo-II intensity is defined as the Myo-II intensity reduced from the intensity at the onset of manipulation. Kymographs are made from the images with KymoResliceWide, after photobleaching correction and registration to remove tissue drift have been performed (all procedures were performed in Fiji).

### Vertex based models

#### Parameter values

We estimated the area of *Drosophila* epithelial cells in our experiments as *A* = 37.4±2.0 μm^2^. In simulation, we obtain a distribution of cell areas close to such value by considering a target area *A*_0_ = 36 μm^2^, which leads to the length scale for our simulation system as 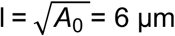. Considering an area stiffness *K*_A_ = 10^6^ N·m^−3^, our stress scale is *σ* = *K*_A_*A*_0_ = 36 pN·μm^−1^. We then normalize Eq. (1) using the length scale 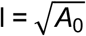, the time scale τ = *γ*/(*K*_A_*A*_0_), and the stress scale *σ*= *K*_A_*A*_0_. By comparing the time evolution of junction length between experiments and simulations, we assume the time scale as τ = *γ*/(*K*_A_*A*_0_) = 1 s. The simulation time step is 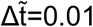. A previous experimental found that the range of tension is 44±11 pN (23). To mimic such heterogeneity of the junction tension we assume a Gaussian distribution of 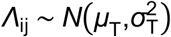 with an average *μ*_T_ = 44 pN 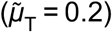 and standard deviation *σ*_T_ = 11 pN 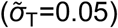. We further validate our choice of the parameter set by finding a quantitative agreement between experiments and simulations on the distribution of cell area, cell perimeter and cell shape index (see Supplementary Information).

#### Trap forces

We randomly choose a junction whose length was within the range 0.9 μm to 2.4 μm (as in experiments), i.e. 0.15 to 0.4 in dimensionless units; this junction will correspond to the junction called middle junction in experiments. The stiffness of the spring connecting the optical trap and the vertex under pulling/pushing is taken as *K*_trap_ = 50 pN·μm^−1^ (23) which results in 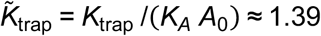. The displacement of the optical trap in experiments is 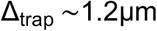, i.e. 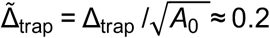.

#### Initialization

We begin with a hexagonal cell pattern consisting of N ≈ 100 cells in a periodic box [0,L_x_]×[0,L_y_], satisfying L_x_L_y_ = N*A*_0_, i.e. <*A*_J_ >_J_ = *A*_0_. For the given distribution in tension 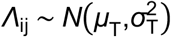, the system is not at steady state; we then let the system relax according to the motion Equation (1). We performed T1 topological transition any time a junction length reaches the threshold value 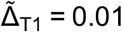 (i.e. Δ_T1_ = 0.06 μm).

Additional information and model validation are provided in the Supplementary Information.

## Movie captions

**Movie 1. Diagonal pull manipulation**

A movie of diagonal pull manipulation which is corresponding to Fig. 1B in the main text (E-cad::GFP). The length change at 60 s post-manipulation is ΔI_60s_ = 1.13 μm. Total duration is 70 s (10 times fast forward). Red circles are the two positions of laser focus. Scale bar: 5 μm.

**Movie 2. Simulation of diagonal pull manipulation**

A movie of numerical simulation of a diagonal pull manipulation using model D as described in the main text.

**Movie 3. Fast shrinkage in direct push manipulation**

A movie of direct push manipulation which is corresponding to Fig. 3B (top) in the main text (E-cad::GFP). The junction was shrunk fast (7 s) after the application of optical forces. Scale bar: 5 μm.

**Movie 4. Slow shrinkage in *direct push* manipulation**

A movie of *direct push* manipulation which is corresponding to Fig. 3B (bottom) in the main text (E-cad::GFP). The junction was slowly shrunk (109 s) after force application. Scale bar: 5 μm.

**Movie 5. Direct push manipulation for a Rock-inhibited tissue**

A movie of *direct push* manipulation for a Rock-inhibited tissue which is corresponding to Fig. 4A in the main text (E-cad::GFP). The junction was not shrunk and escaped from optical manipulation. Scale bar: 5 μm.

