## Supplementary Information for "Two-point optical manipulation reveals mechanosensitive remodeling of cell-cell contacts in vivo"

(Dated: July 19, 2022)

**Contents**

**I. Experimental methods** 1
A. Dextran injection 1
B. Changes in junction length after trap release 2

**II. Model methods** 3
A. Vertex model motion equation 3
B. Junctional tension/strain remodeling 4
C. Myosin accumulation induced by junction shrinkage 4
D. Estimation of the parameter values 5
E. Numerical procedure 5

**References** 8

### I. EXPERIMENTAL METHODS

#### A. Dextran injection

To detect potential large-size disruption of the junctions or changes in the axial position of cells during manipulation, we injected fluorescently labeled Dextran (10000 MW, -Alexa568, Cie) between the vitelline membrane and the apical surface of the epithelial tissue (Fig. S1). Dextran cannot enter cells but diffuses fast between them, thus marking the extracellular space. The laser spots are located at the adherens junction plane located at 1µm distance from the apical surface. We did not detect any significant changes of Dextran signal at junctions during manipulation (Fig. S1 B). This observation rules out the possibility that changes in junction length are due to damage of the junctions or axial movement of the cells due to optical force application.

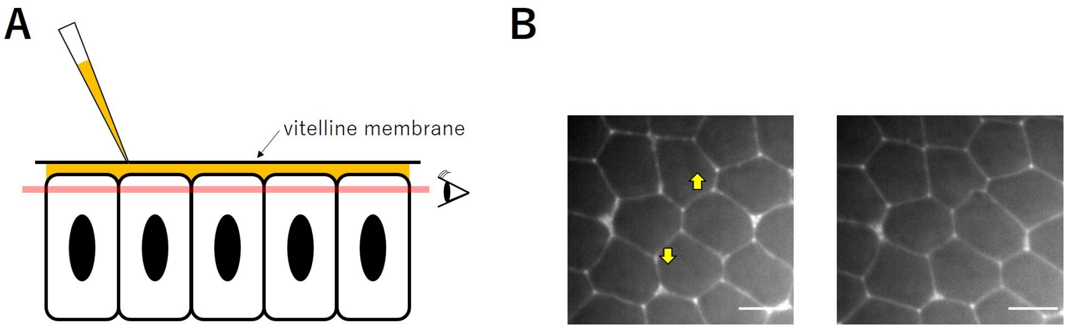


Figure S1: (A) Fluorescently-labeled dextran solution is injected the perivitelline space. (B) The traps are moved away from each other, in antiparallel direction and maintained at 1 µm distance from their initial position (yellow arrows) causing junction remodeling. The right image shows the epithelial cells 1 minute after the onset of manipulation. Scale bar: 5 µm.

#### B. Changes in junction length after trap release

Tension dynamical remodeling manifests in the adaptation of tension to strains. Such adaptation is encapsulated in the rates of tension remodeling under contraction or extension, kc and ke respectively. A prediction of negative (or positive) kc is that after a junction contraction, tension decreases (or increases). To verify this prediction, we performed direct push manipulation in conditions where Myosin-II mechanosensitive response was abolished (Rock inhibited tissues). In such conditions, direct push manipulation produces only partial shrinkage, and the vertices escape the optical traps short after manipulation (Fig. 4A). We measured the junction length after trap release and at the onset of manipulation (Fig. S2). Junction length was always found larger after trap release than at the onset of manipulation. Such extension suggests that the tension of the junction is decreased after contractile strain, which is consistent with $k_c < 0$.

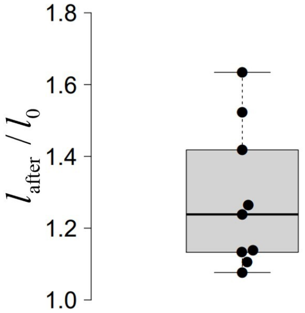


Figure S2: Ratio between the initial middle junction length and its length after direct push manipulation.

### II. MODEL METHODS

#### A. Vertex model motion equation

We use the vertex model to simulate *Drosophila* epithelium under local pulling/pushing forces applied by optical tweezers. In the vertex model, a epithelium cell sheet is described by a two-dimensional planar polygonal network where cells are represented by interconnected polygons. Cell–cell interfaces are traditionally characterized by straight lines which connect two neighboring tricellular junctions. Here, for the sake of mimicking pulling/pushing forces applied at the middle of some cell–cell interfaces, we divide each cell–cell interface into two segments, as shown in Fig. S3.

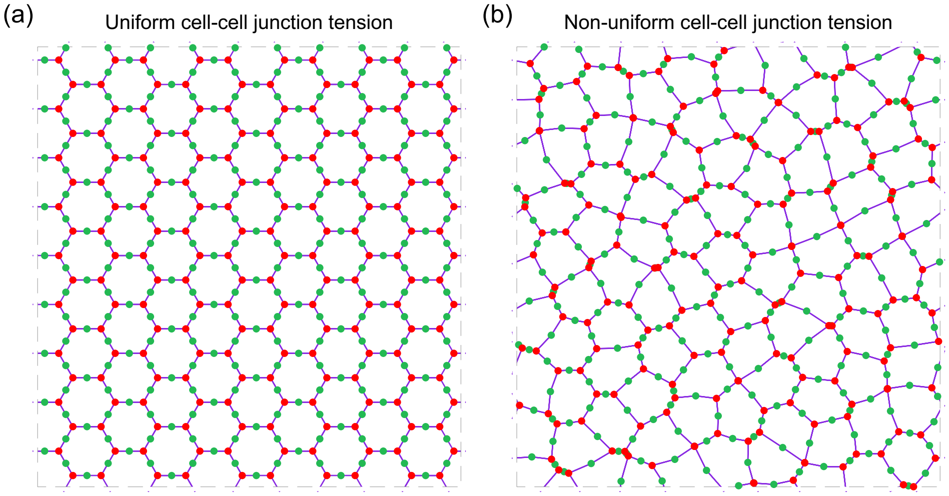


Figure S3: Schematic of an epithelial vertex model. Shown here are two typical cell sheet patterns with different variation $\sigma_T$ of cell-cell junction tension. Tricellular junctions are marked as red; while middle vertices are marked as green. (a) Uniform cell–cell junction tension with $\tilde{\sigma}_T = 0$. (b) Non-uniform cell–cell junction tension with $\tilde{\sigma}_T = 0.05$. For other parameter values, see Table I.

The dynamics of the epithelium is dictated by the force balance equation at each vertex $i$. Assuming pure frictional dynamics, the force balance equation reads:

$$\underbrace{\mathbf{F}_i^{(\text{friction})}}_{\text{friction force}} + \underbrace{\mathbf{F}_i^{(\text{passive})}}_{\text{internal passive force}} + \underbrace{\mathbf{F}_i^{(\text{trap})}}_{\text{external pulling/pushing force}} = \mathbf{0}, \tag{S1}$$

which accounts for a balance between the friction forces, the internal passive forces, and the external pulling/pushing forces. In details, the first term represents for the friction force between cells and the environment, which can be simply expressed as, $\mathbf{F}_i^{(\text{friction})} = -\gamma \mathbf{v}_i$, with $\gamma$ being the friction coefficient and $\mathbf{v}_i = \mathrm{d}\mathbf{r}_i/\mathrm{d}t$ being the velocity of vertex $i$. The second term is the internal passive forces stemming from a mechanical energy $E$ such that $\mathbf{F}_i^{(\text{passive})} = -\partial E/\partial \mathbf{r}_i$; while the third term refers to the external pulling/pushing force applied by the optical tweezers.

*Trap force* – To mimic the locally applied forces, we assume a elastic spring between the optical trap and the vertex under pulling/pushing. Assuming linear elasticity, the external pulling/pushing force can be written as

$$\mathbf{F}_i^{(\text{trap})} = \begin{cases} K_{\text{trap}} \left(\mathbf{r}_{i,\text{trap}} - \mathbf{r}_i\right) & , \quad \text{if vertex } i \text{ is under pulling/pushing} \\ \mathbf{0} & , \quad \text{otherwise} \end{cases}, \tag{S2}$$

where $K_{\text{trap}}$ is the stiffness of the elastic spring connecting the optical trap and the vertex under pulling/pushing; $\mathbf{r}_{i,\text{trap}}$ is the position of the optical trap that pull or push vertex $i$.

*Mechanical energy* – We express the mechanical energy of a *Drosophila* epithelium as [1–4],

$$E = \underbrace{\sum_J \frac{1}{2} K_A (A_J - A_0)^2}_{\text{area elasticity}} + \underbrace{\sum_{\langle i,j \rangle} \Lambda_{ij} l_{ij}}_{\text{interfacial tension}}, \tag{S3}$$

where the two terms account for the cell area elasticity and the cell–cell interfacial tension, respectively. In details, $K_A$ is the area stiffness of cells; $A_J$ is the area of the $J-$th cell and $A_0$ is the preferred area. In addition, $\Lambda_{ij}$ represents the dynamic interfacial tension (which remodels dynamically according to junction strain, see Eq. (S4)) of the junction $ij$, and $l_{ij} = |\mathbf{r}_i - \mathbf{r}_j|$ is the junction length.

To mimic the heterogeneity of the cell–cell interfacial tension $T_{ij}$, before pulling/pushing perturbations ($t < 0$), we assume a Gaussian distribution of $\Lambda_{ij}$, $\Lambda_{ij} \sim \mathcal{N}(\mu_T, \sigma_T^2)$ with $\mu_T$ being the mean value and $\sigma_T$ being the standard deviation.

*Normalization* – In simulations, we normalize the above governing equations using the length scale $\ell = \sqrt{A_0}$, the time scale $\tau = \gamma/(K_A A_0)$, and the stress scale $\sigma = K_A A_0$.

#### B. Junctional tension/strain remodeling

Previous study suggests a strain-dependent tension remodeling behavior of cell–cell junctions [5]. Specifically, the cell–cell junction tension remodelling can be described by the following dynamic equation,

$$\frac{\mathrm{d}\Lambda_{ij}}{\mathrm{d}t} = \begin{cases} -k_e\left(l_{ij} - l_{0,ij}\right) & \varepsilon_{ij} > \varepsilon_{cr} \\ 0 & -\varepsilon_{cr} < \varepsilon_{ij} < \varepsilon_{cr} \\ -k_c\left(l_{ij} - l_{0,jj}\right) & \varepsilon_{ij} < -\varepsilon_{cr} \end{cases} \tag{S4}$$

where $k_e$ and $k_c$ are the tension remodeling rates under junction extension or contraction, respectively; $\varepsilon_{ij} = (l_{ij} - l_{0,ij})/l_{0,ij}$ is the strain of the cell–cell junction $ij$ with $l_{0,ij}$ being the rest length; $\varepsilon_{cr}$ is a critical strain beyond which cell–cell junction tension remodeling is triggered.

Further assuming continuous strain relaxation of cell–cell junctions [5], the rest length $l_{0,ij}$ of the cell–cell junction $ij$ evolves according to

$$\frac{1}{l_{0,ij}}\frac{\mathrm{d}l_{0,ij}}{\mathrm{d}t} = k_L \varepsilon_{ij}, \tag{S5}$$

where $k_L > 0$ is the relaxation rate of rest cell–cell junction length $l_{0,ij}$ to the current cell–cell junction length $l_{ij}$. At steady state, we have $l_{ij} = l_{0,ij}$.

#### C. Myosin accumulation induced by junction shrinkage

*Strain-rate tension modulation* – Our experiments reveal that junction shrinkage can induce myosin accumulation at cell–cell junctions that undergo large shrinking rates ($\dot{\varepsilon}_{ij} < \dot{\varepsilon}_{\mathrm{cr}}^{(\mathrm{myo})} < 0$). Here, we define the final model (called model D in the main text) where the myosin accumulation induced tension $T_{ij}^{(\mathrm{myo})}$ evolves according to the junction strain rate $\dot{\varepsilon}_{ij}(t)$ according to the relation

$$T_{ij}^{(\mathrm{myo})}(t) = \int_{-\infty}^{t} H\left[\dot{\varepsilon}_{cr}^{(\mathrm{myo})} - \dot{\varepsilon}_{ij}(\tau)\right]\frac{T_m}{\tau_m}\exp\left(-\frac{t-\tau}{\tau_m}\right)\mathrm{d}\tau, \tag{S6}$$

where $\dot{\varepsilon}_{cr}^{(\mathrm{myo})} < 0$ is a strain rate threshold below which there will be myosin accumulation at the junction $ij$; the junction strain rate $\dot{\varepsilon}_{ij}$ is defined as $\dot{\varepsilon}_{ij} = \mathrm{d}\varepsilon_{ij}/\mathrm{d}t$; $T_m$ is the tension magnitude induced by myosin accumulation; $\tau_m$ is a time scale of the tension increase induced by myosin accumulation.

*Numerical scheme* – The myosin accumulation induced tension $T_{ij}^{(\mathrm{myo})}(t)$ defined by Eq. (S6) can be numerically implemented as below:

$$T_{ij}^{(\mathrm{myo})}(t+\Delta t) = T_{ij}^{(\mathrm{myo})}(t)\exp\left(-\frac{\Delta t}{\tau_m}\right) + H\left[\dot{\varepsilon}_{cr}^{(\mathrm{myo})} - \dot{\varepsilon}_{ij}(t+\Delta t)\right]T_m\left[1-\exp\left(-\frac{\Delta t}{\tau_m}\right)\right]. \tag{S7}$$

We give the proof as below. Letting

$$f(t,\tau) = H\left[\dot{\varepsilon}_{cr}^{(\mathrm{myo})} - \dot{\varepsilon}_{ij}(\tau)\right]\frac{T_m}{\tau_m}\exp\left(-\frac{t-\tau}{\tau_m}\right), \tag{S8}$$

$T_{ij}^{(\text{myo})}(t)$ can be expressed as,

$$T_{ij}^{(\text{myo})} = \int_{-\infty}^{t} f(t,\tau)\,\mathrm{d}\tau. \tag{S9}$$

Therefore, we have the updating scheme of $T_{ij}^{(\text{myo})}(t)$ as,

$$\begin{aligned} T_{ij}^{(\text{myo})}(t+\Delta t) &= \int_{-\infty}^{t+\Delta t} f(t+\Delta t,\tau)\,\mathrm{d}\tau = \int_{-\infty}^{t} f(t+\Delta t,\tau)\,\mathrm{d}\tau + \int_{t}^{t+\Delta t} f(t+\Delta t,\tau)\,\mathrm{d}\tau \\ &= \exp\left(-\frac{\Delta t}{\tau_m}\right)\int_{-\infty}^{t} f(t,\tau)\,\mathrm{d}\tau + \int_{t}^{t+\Delta t} f(t+\Delta t,\tau)\,\mathrm{d}\tau \\ &\approx T_{ij}^{(\text{myo})}(t)\exp\left(-\frac{\Delta t}{\tau_m}\right) + H\left[\dot{\varepsilon}_{cr}^{(\text{myo})} - \dot{\varepsilon}_{ij}(t+\Delta t)\right] T_m \left[1-\exp\left(-\frac{\Delta t}{\tau_m}\right)\right], \end{aligned} \tag{S10}$$

where we have used the formula $f(t+\Delta t,\tau) = f(t,\tau)\exp(-\Delta t/\tau_m)$.

### D. Estimation of the parameter values

*Area forces* We measured the area of *Drosophila* epithelial cells as $A = 37.4 \pm 2.0\ \mu\text{m}^2$ (see Fig. S4(a)). Thus assuming the incompressibility of cells, we can roughly take $A_0 = 36\ \mu\text{m}^2$, which leads to the length scale for our simulation system as $\ell = \sqrt{A_0} = 6\ \mu$m. Assuming the cell area stiffness $K_A = 10^6\ \text{N}\cdot\text{m}^{-3}$ [6], we then have the stress scale as $\sigma = K_A A_0 = 36\ \text{pN}\cdot\mu\text{m}^{-1}$.

*Tension forces* Previous experimental study measured that the tension $T_{ij}$ is on the order of $44 \pm 22$ pN [7]. We thus set the mean cell–cell interfacial tension as $\mu_T = \langle T_{ij}\rangle = 44$ pN.

*Trap forces* The stiffness of the spring connecting the optical trap and the vertex under pulling/pushing is taken as $K_{\text{trap}} = 50\ \text{pN}\cdot\mu\text{m}^{-1}$ [7], which results in $\tilde{K}_{\text{trap}} = K_{\text{trap}}/(K_A A_0) \approx 1.39$. The displacement of the optical trap in experiments is $\Delta_{\text{trap}} \sim 1.2\ \mu$m, i.e. $\tilde{\Delta}_{\text{trap}} = \Delta_{\text{trap}}/\sqrt{A_0} \approx 0.2$.

*Friction* By comparing the time evolution of junction length between experiments and simulations, we assume the time scale as $\tau = \gamma/(K_A A_0) = 1$ s.

Therefore, in our simulations, we set the values of dimensionless parameters as below: $\tilde{\mu}_T = 0.2$, $\tilde{\sigma}_T = 0.05$, $\tilde{K}_{\text{trap}} = 1.4$, $\tilde{\Delta}_{\text{trap}} = 0.2$, and $\Delta\tilde{t} = 0.01$ (see also Table I).

Using such a parameter set, we have obtained a cellular pattern as shown in Fig. S3(b). We further compute the distribution of cell area, cell perimeter and cell shape index, and compare them with experiments, as shown in Fig. S4. It demonstrates that for the epithelial pattern obtained from our simulations the distributions of geometric properties of cells are similar to those of our experiments.

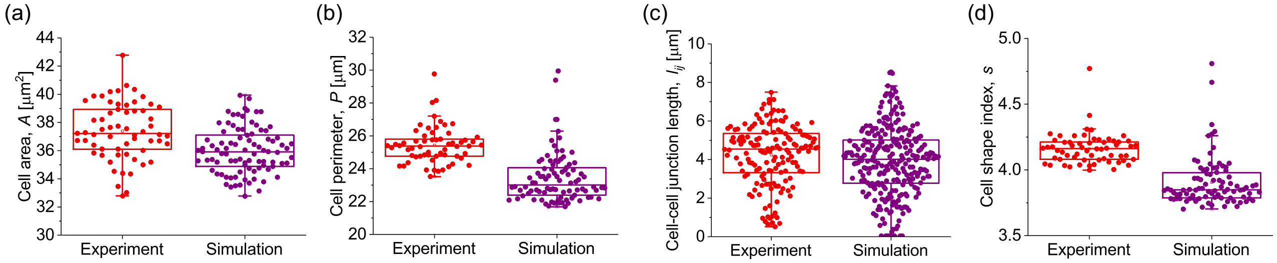


Figure S4: Two-by-two comparison of cellular configuration distribution in experiments (red) and simulations (magenta): (a) cell area $A$; (b) cell perimeter $P$; (c) cell–cell junction length $l_{ij}$; (d) cell shape index $s = P/\sqrt{A}$. Parameters defined Table I.

### E. Numerical procedure

Here we briefly give the simulation procedure as below:

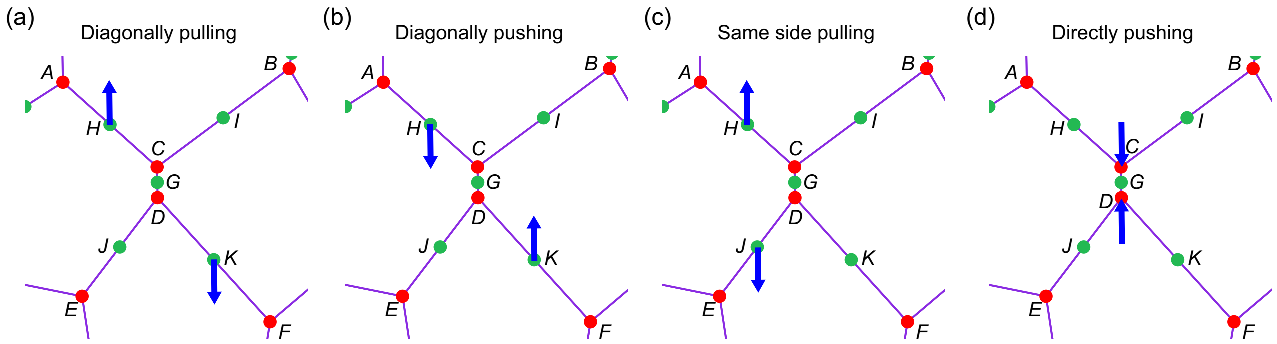


Figure S5: Schematic of applying locally pulling/pushing forces. (a) Diagonally pulling case, where the pulling forces are applied at vertices $H$ and $K$ in opposite directions (parallel to the junction $C-D$ initially) according to our optical trap model, see Eq. (S2). (b) Diagonally pushing case, where the pushing forces are applied at vertices $H$ and $K$ in opposite directions (parallel to the junction $C-D$ initially). (c) Same side pulling case, where the pulling forces are applied at vertices $H$ and $J$ in opposite directions (parallel to the junction $C-D$ initially). (d) Directly pushing case, where the pushing forces are applied at vertices $C$ and $D$ in opposite directions (parallel to the junction $C-D$ initially).

1. We begin with a hexagonal cell pattern (see Fig. S3(a)) consisting of $N \approx 100$ cells in a periodic box $[0, L_x] \times [0, L_y]$, satisfying $L_x L_y = N A_0$, i.e. $< A_J >_J = A_0$. We then randomly set the tension $\Lambda_{ij}$ of each cell–cell junction $ij$, satisfying a Gaussian distribution, i.e. $\Lambda_{ij} \sim \mathcal{N}(\mu_T, \sigma_T^2)$. The system is relaxed to a steady state (see Fig. S3(b)), according to the motion equation (S1) (mind that, at this stage, $\mathbf{F}_i^{(\mathrm{trap})} = \mathbf{0}$ here). At each simulation time step, we estimate the new vertex position through a forward Euler scheme as $\mathbf{r}_i(t+\Delta t) = \mathbf{r}_i(t) + \mathbf{v}_i \Delta t$. T1 topological transition are implemented any time the length threshold reaches the value $\tilde{\Delta}_{T1} = 0.01$ (i.e. $\Delta_{T1} = 0.06\ \mu$m).

2. Once we obtain the relaxed cell configuration (Fig. S3(b)), we randomly choose a short junction (with the length $l_{ij}$ satisfying $0.9\ \mu\mathrm{m} < l_{ij} < 2.4\ \mu\mathrm{m}$, i.e. $0.15 < \tilde{l}_{ij} < 0.4$) of the tissue, which will correspond to the junction called middle junction in experiments (see the junction $C-D$ in Fig. S5).

3. We then add at $t = 0$ pulling/pushing forces to the vertices corresponding to the experimental setup of interest (see Fig. S5 for details). Specifically, we set the position of the associated optical trap acting on a vertex $i$ as $\mathbf{r}_{i,\mathrm{trap}}(t=0) = \mathbf{r}_i(t=0) + \Delta_{\mathrm{trap}} \mathbf{t}_{i,\mathrm{trap}}$ with $\Delta_{\mathrm{trap}}$ being the displacement magnitude of the trap and $\mathbf{t}_{i,\mathrm{trap}}$ being the pulling/pushing direction. We then fix the position of the optical trap $\mathbf{r}_{i,\mathrm{trap}}(t) = $ constant and let the system evolve according to the motion equation (S1).

TABLE I: List of parameter values used in simulations.

| Parameter | Description | Value | Dimensionless value |
|---|---|---|---|
| $K_A$ | Cell area stiffness | $10^6$ N $\cdot$ m$^{-3}$ [6] | 1 |
| $A_0$ | Preferred cell area | 36 $\mu$m$^2$ | 1 |
| $\mu_T$ | Average cell junctional tension | 44 pN [7] | 0.2 |
| $\sigma_T$ | Standard derivation of cell junctional tension | 11 pN | 0.05 |
| $\Delta_{\mathrm{trap}}$ | Displacement of the optical trap | 1.2 $\mu$m | 0.2 |
| $K_{\mathrm{trap}}$ | Stiffness of the optical trap | 50 pN $\cdot$ $\mu$m$^{-1}$ [7] | 1.4 |
| $k_L$ | Remodeling rate of rest length | 0.04 s$^{-1}$ | 0.04 |
| $k_e$ | Tension remodeling rate under extension | 0.18 pN $\cdot$ $\mu$m$^{-1}$ $\cdot$ s$^{-1}$ | 0.005 |
| $k_c$ | Tension remodeling rate under contraction | $-0.108$ pN$\cdot\mu$m$^{-1}\cdot$s$^{-1}$ | $-0.003$ |
| $\varepsilon_{cr}$ | Strain threshold for tension remodeling | 0.1 | 0.1 |
| $\dot{\varepsilon}_{cr}^{(\mathrm{myo})}$ | Strain rate threshold for myosin recruitment at cell–cell junctions | $-0.05$ s$^{-1}$ | $-0.05$ |
| $T_m$ | Tension magnitude induced by myosin recruitment at cell–cell junctions | 54 pN | 0.25 |
| $\tau_m$ | Time scale of myosin recruitment at cell–cell junctions | 10 s | 10 |
| $\Delta t$ | Simulation time step | 0.01 s | 0.01 |
| $\ell = \sqrt{A_0}$ | Length scale | 6 $\mu$m | 1 |
| $\tau = \gamma/(K_A A_0)$ | Time scale | 1 s | 1 |
| $\sigma = K_A A_0$ | Stress scale | 36 pN $\cdot$ $\mu$m$^{-1}$ | 1 |
